# A whole-genome sequenced control population in northern Sweden reveals subregional genetic differences

**DOI:** 10.1101/2020.02.18.933622

**Authors:** Daniel Svensson, Matilda Rentoft, Anna M. Dahlin, Emma Lundholm, Pall I. Olason, Andreas Sjödin, Carin Nylander, Beatrice S. Melin, Johan Trygg, Erik Johansson

## Abstract

The number of national reference populations that are whole-genome sequenced are rapidly increasing. Partly driving this development is the fact that genetic disease studies benefit from knowing the genetic variation typical for the geographical area of interest. A whole-genome sequenced Swedish national reference population (n=1000) has been recently published but with few samples from northern Sweden. In the present study we have whole-genome sequenced a control population (n=300) (ACpop) from Västerbotten County, a sparsely populated region in northern Sweden previously shown to be genetically different from southern Sweden. The aggregated variant frequencies within ACpop are publicly available (DOI 10.17044/NBIS/G000005) to function as a basic resource in clinical genetics and for genetic studies. Our analysis of ACpop, representing approximately 0.11% of the population in Västerbotten, indicates the presence of a genetic substructure within the county. Furthermore, a demographic analysis showed that the population from which samples were drawn was to a large extent geographically stationary, a finding that was corroborated in the genetic analysis down to the level of municipalities. Including ACpop in the reference population when imputing unknown variants in a Västerbotten cohort resulted in a strong increase in the number of high-confidence imputed variants (up to 81% for variants with minor allele frequency < 5%). ACpop was initially designed for cancer disease studies, but the genetic structure within the cohort will be of general interest for all genetic disease studies in northern Sweden.

## Introduction

The challenge for all studies on genetic diseases is to disentangle disease-causing genetic variations from those that are not. This task is inherently dependent on knowledge of the genetic variation in the population. Even though a number of large-scale projects (1–3) have mapped much of the common global human genetic variation, the map of human genetic diversity is far from complete, in particular with respect to rare variants. The recent explosive population growth has resulted in an abundance of rare variants (4,5) which, because of their relatively recent origin (6,7) tend to be geographically clustered to a higher degree than variants that are common in different parts of the world (8). Therefore, in order to obtain a better overview of all genetic variation it is important to sequence geographically focused populations. In recent years, several such projects have mapped the genetic variation in national populations, providing important information regarding locally occurring rare variation (9–11).

An addition to these was the recent release of a whole-genome sequenced Swedish national reference population, SweGen (12). The genetic structure of the Swedish population is characterized by a pronounced genetic difference between the northernmost counties and the rest of Sweden (13,14). For example, the genetic distance, as measured by F_ST_, between the northern and southern parts of Sweden is larger than that between southern Sweden and other populations of Northern European descent (13). Furthermore, the northernmost counties exhibit an elevated level of homozygosity (13,14) and an increase in the genetic distance as a function of the geographical distance (14), compared to southern Sweden. In agreement with these observations, Y chromosomal haplotype frequencies also showed that the population in Västerbotten County in northern Sweden is differentiated from that in the southern Swedish counties (15).

Our studies focus on this genetically pronounced northern Swedish region, specifically Västerbotten County. The county is characterized by vast distances between population centers; the geographical footprint is large (54,665 km^2^) (16), but the population size is small (268,278) (17). The demographic history of the county highlights a sparsely populated region with an estimated population of as few as 7,304 individuals in 1571 who mainly inhabited the coastal region (18). Historically, the inland region has predominantly been inhabited by the indigenous Sami population, but settlement programs were initiated by the state during the mid- to late-17^th^ century with the main part of the in-migration taking place from mid-18^th^ to mid-19^th^ century (19). It was also in this later period that settlers formed the majority in relation to the Sami population (20). Previous work based on marriage patterns and distribution of allele frequencies of eight protein markers indicated the presence of a genetic substructure within Västerbotten County (21). The hypothesized substructure was associated with the major river valleys that stretch from the mountains in the west to the coast in the east in northern Sweden.

In summary, the genetics of the Västerbotten population have been shown to be distinct compared to the rest of Sweden. To improve the likelihood of conducting successful studies on genetic disease in this region, there is a need to acquire a better understanding of the genetic variation in the population. While focusing on establishing a control population to be used in cancer disease studies, we have whole-genome sequenced a population sample named ACpop from the county of Västerbotten. The whole-genome sequenced population, ACpop, provides a high-resolution map of the genetic variation which agrees with earlier studies of protein markers and marriage patterns. Imputation of rare variants in genotyping data from the region is also evaluated.

## Materials and methods

### Sample selection

The study was approved by the ethics board in Umeå, Sweden (dnr 2014-290-31 and dnr 2017-370-32). The 300 samples were selected from the Västerbotten Intervention Programme (22) (VIP) cohort of the Northern Sweden Health and Disease Study (NSHDS). Blood from residents were collected in health care units in each municipality and a donated sample was considered to represent a resident of that municipality. Half of the samples to be sequenced were spread evenly over the 15 municipalities of the county. The remaining half were dedicated to increase the frequency of sequenced individuals in the three most highly populated municipalities (Lycksele, Skellefteå, and Umeå), as well as two municipalities that is located on the border (Supplementary Fig 1). An equal number of men and women were selected in each municipality. To maximize the diversity among selected individuals and to minimize selection bias, 27 phenotypic, health, and lifestyle-related variables were extracted from the VIP (Supplementary Table 1), and a principal component (PC) model was used to select the individuals to be sequenced from each municipality. Principal component (PC) models were calculated separately for each gender and municipality. Samples to be sequenced were selected from the two first PCs according to a full factorial design in two levels with one center point (Supplementary Fig 2). For municipalities where sample size was substantially increased (Lycksele, Umeå, and Skellefteå), we extended the experimental design with another full factorial design around each of the corner points, as well as with another four center points (Supplementary Fig 2b). For the municipalities where sample size was moderately increased (Storuman and Malå), we extended the experimental design simply by adding another full factorial design in two levels and one center point but angled 45 degrees to the original one.

### Whole genome sequencing, raw data processing and calling of variants

DNA samples were obtained from the NSHDS and paired-end sequenced (2×150 bp) to a depth of at least 30x using the Illumina HiSeqX system at NGI-U (Uppsala, Sweden). Samples underwent a quality check by the sequencing facility. PCR-free library preparation kits were used for all samples.

Raw data processing was performed in accordance with the GATK best practices(23,24). Briefly, reads were aligned to the 1000g fasta reference (b37) using BWA (v0.7.10-r789) (25) Sorting, indexing, and marking of duplicates was done using Picard (v.1.118) (26), and realignment around indels was done using GATK (v3.3.0) (27). Qualimap (v2.0.2) (28) was used to assure sample quality and to identify any deviating samples. SNPs and small indels were called for each sample separately using HaplotypeCaller (GATK v.3.3.0). The resulting gVCFs were then jointly genotyped into a single VCF file using GenotypeGVCFs (GATK v. 3.3.0). The called variants were quality filtered in accordance with GATK recommendations using VQSR. Guided by the Ti/Tv ratio, the truth sensitivity cutoff was set to 99.7 and 99.0 for SNPs and small indels, respectively. Additionally, variants were annotated with the result from a Hardy-Weinberg equilibrium (HWE) test. Variants with a Phred-scaled p-value above 60 (p ≤ 10^−6^) were removed in all analyses.

### Cryptic relations analysis

The software KING (29) was used to screen for unknown pairwise relationships between all samples in ACpop. KING was run with the *kinship* flag and using all biallelic SNPs with a successful genotyping rate of at least 99.9%. Pairwise relationships were inferred using the kinship coefficient as suggested by (29).

### PCA

Both PCA models were produced using smartpca (30) (v. 13050) of the EIGENSOFT (v. 6.0.1) software package. The allowed maximum missing genotype rate was 0.1% and 1% for variants and individuals, respectively. Two full sibling relations (4 individuals) were suggested to exist by the relationship analysis, and one individual from each of these relations was excluded prior to PCA. For the joint PCA of ACpop and the 1000 genomes (1000g) project European super population, autosomal biallelic SNPs with a frequency of at least 5% in ACpop and each European subpopulation were used. For the PCA of only ACpop, autosomal biallelic SNPs with a frequency of at least 5% were used. Variants overlapping long-range LD regions were excluded as suggested by EIGENSOFT. A linkage disequilibrium (LD) pruning step was performed in both cases, both using PLINK (31) (v1.90b3) with a window size of 20,000, step size of 2,000 and an r^2^ threshold of 0.2. In total, 118,544 variants were used for the joint PCA and 156,680 variants for the ACpop PCA. The 1000g genotypes (release v5 20130502) were downloaded from ftp://ftp.1000genomes.ebi.ac.uk/vol1/ftp/release/20130502/.

### Imputation

Imputation was performed in two different cohorts. One cohort of individuals from Västerbotten county (n = 500) and one cohort of Swedish individuals (n = 214). Both cohorts have previously been used as control populations in a GWAS of glioma (32), and were genotyped with the Illumina 660W array. All individuals had a call rate >94% and were unrelated (PI-HAT< 0.2). SNPs were filtered based on call rate (>90%), minor allele frequency (>0.01), and HWE (p>1×10^−6^). A/T and G/C SNPs were removed. All subjects were phased together using SHAPEIT (33) and untyped genotypes on chromosome 20 were imputed with IMPUTE2 (34,35) (v2.3.2) using three different reference populations – 1000g, ACpop, and combination of the two (1000g + ACpop). Imputation was performed on genomic regions less than 5 Mb in size with 1 Mb buffer regions. To create the ACpop reference panel we first created a VCF by jointly calling variants in the ACpop samples and in 64 whole genome sequenced individuals from two families. The families originate from the region. Raw data processing and calling of variants was performed in the same way as was described for ACpop. Phasing was performed with SHAPEIT2 (36), using all biallelic SNPs with a genotyping rate above 1%. SHAPEIT2 was run with the duohmm option active. The number of conditioning states was set to 1000, the window size was set to 300, and the MCMC iterations were set to have 15 burn-in and pruning iterations and 50 main iterations. After phasing, we extracted the haplotypes from the 300 ACpop samples and used them for subsequent imputation analysis. The 1000g reference panel was downloaded from https://mathgen.stats.ox.ac.uk/impute/1000GP_Phase3.html and is based on 2,504 samples from phase 3 (v5 20130502).

### F_ST_

Pairwise F_ST_ was calculated using smartpca (30) (v. 13050) from the EIGENSOFT (v. 6.0.1) software package. The same two sibling samples that were excluded in the PCA were also excluded in this analysis. Variants with a missing genotype rate of 0.1% or more were excluded from the analysis. For each pairwise calculation, only autosomal biallelic SNPs that were polymorphic (alternative allele frequency > 0) in both investigated populations were used. Pairwise F_ST_ was calculated between ACpop and the European subpopulations of the 1000g project (release v5 20130502) as well as between regionally defined populations within ACpop.

### Inbreeding analysis

Calculation of the inbreeding coefficient for ACpop was performed using FSuite (37) (v. 1.0.4). We followed the procedure outlined in Gazal et al (38) and included the 26 populations of the 1000g project in the analysis to corroborate the results (data not shown). We included autosomal biallelic SNPs with a frequency of at least 5% in each of the 27 populations and which did not deviate from HWE (p < 10^−5^) in any of the 27 populations. The HWE test was performed using PLINK (31,39) (v1.90b3). Additionally, we removed variants with a missing genotype rate of 5% or more, as suggested by FSuite manual. Prior to calculating the population specific allele frequencies, 14 individuals from 15 reported (38) first-cousin or closer relationships (including one trio) were excluded from the 1000g data (release v5 20130502), as well as the same two ACpop individuals that were excluded in the PCA and F_ST_ analyses due to close relationships. FSuite relies on the creation of several random sparse genome maps (submaps) in order to avoid markers in linkage disequilibrium (40). We selected markers between recombination hotspots for a total of 100 submaps. The inbreeding coefficient reported by FSuite is the median inbreeding coefficient from the 100 submaps.

## Results

### Selection of the cancer control samples results in a population that reflects diverse Västerbotten origins

We have whole genome sequenced 300 individuals intended to be used as a control population in genetic disease studies in the Swedish county of Västerbotten and northern Sweden. Based on a previous study (21), it was clear that the control population would benefit from a selection of individuals from all parts of Västerbotten County. The longitudinal and large-scale health study cohort, the Västerbotten Intervention Programme (22) (VIP) cohort of the Northern Sweden Health and Disease Study (NSHDS), provided approximately 95,000 samples from which a selection could be made. Through the VIP, individuals across Västerbotten were invited to undergo a health examination, complete a questionnaire on health and lifestyle, and donate a blood sample for research. Furthermore, our primary focus was to use the control population for cancer studies, thus only individuals who had reached at least 80 years of age in 2014 without being diagnosed with cancer were included from the VIP cohort. This resulted in 2117 women and 1431 men from which the final selection of 150 women and 150 men was made (Figure 1A). The included individuals had donated their blood samples at the latest in 1995 and were at least 60 years of age when making the donation.

**Figure 1.**
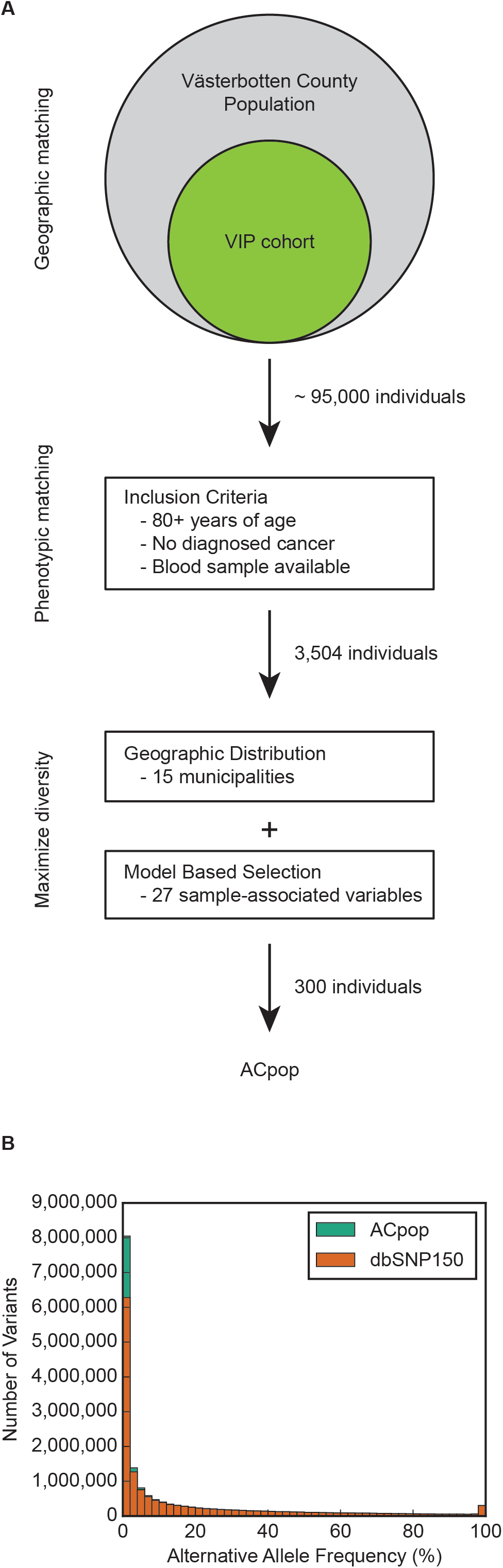
Selection process and allele frequency distribution. The initial aim of ACpop was to sequence a geographically and phenotypically matched cancer control population for Västerbotten County. The selection was made using three steps; selection of cohort from which individuals were identified, inclusion criteria, and maximization of diversity. (**A**). The alternative allele frequency distribution of the variants contained in ACpop, stratified over 50 bins of 2% each. The share of the variants in each bin that does not overlap with dbSNP v. 150 is indicated by green, while the opposite is indicated by orange (**B**).

The municipality of birth records for all residents in Västerbotten County in 1995 who were born 1934 and earlier were utilized to investigate the origins of the population (available in the Linnaeus database, Centre for Demography and Ageing, Umeå University). An analysis of the population demography revealed that 82% of the Västerbotten population aged 60 years or older in 1995 was born in Västerbotten, and 59.4% of the population still resided in the same municipality where they were born (Table 1). Only 10.3% of the population had migrated from a neighboring county, and 7.7% of the population came from other parts of Sweden (further south) or were immigrants. The inflow is expected to increase with successive cohorts. When comparing to a more recent cohort of 60 years and older in 2013 it is found that the share of the population originating from other parts of Sweden and immigrants have risen to 13% and 11% respectively. Mobility into Västerbotten has been low in the 1995 cohort but mobility out from Västerbotten was higher as 35 % of those born in the county lived outside Västerbotten in 1995. Based on the low mobility, the 300 individuals were selected from all 15 municipalities in Västerbotten (Supplementary Figure 1) to obtain a sample of the genetic variation from all parts of the county. A selection strategy was employed for each municipality to maximize diversity and avoid random selection bias. In short, the strategy consisted of using principal component analysis (PCA) to summarize 27 sample-associated features in conjunction with design of experiments (Supplementary table 1).

**Table 1.** Origins of the Västerbotten population aged 60 or older in 1995. The origin (geographical region of birth) as a percentage of the population that were living in Västerbotten County in the year 1995 and that were aged 60 or more at the time.

| **Geographical region of birth** | **Men (%)** | **Women (%)** | **Total (%)** |
| --- | --- | --- | --- |
| **Västerbotten County, same municipality** | 63.8 | 55.9 | 59.4 |
| **Västerbotten County, other municipality** | 20.3 | 24.4 | 22.6 |
| **Neighboring counties** | 9.3 | 11.1 | 10.3 |
| **Non-neighboring counties in Sweden** | 4.7 | 4.4 | 4.5 |
| **Other country** | 1.9 | 4.2 | 3.2 |

### Overview of the ACpop variant dataset

The final call set comprised 17,344,482 variants, distributed over 16,522,463 variant sites, after quality control. Of these variants, 14,513,111 and 2,831,371 were SNPs and indels, respectively. A large share of the variants was present at a low-frequency (Figure 1B). For example, 6,176,006 variants have an allele frequency (AF) of below 1%, of which half (2,941,762) were seen only once (singletons, allele count of 1). As sequencing efforts continue around the world, the total number of discovered genetic variants increases. The most comprehensive collection of genetic variants is dbSNP, which provides updated releases on a regular basis. The latest release of dbSNP (v. 150) contains a total of over 325.7 M variants, more than double the size of the previous release (v. 149). We compared the ACpop variant set against the latest release of dbSNP (v. 150) and found that a total of 2,022,713 variants were not represented in the collection (Table 2). Even though many (938,209) of these unique variants were singletons, as many as 471,744 and 115,682 variants had an AF ≥ 1% and AF ≥ 5%, respectively. Compared to the recently released whole genome sequenced Swedish national reference (SweGen), ACpop contains 1,813,725 unique variants, and a substantial number (794,800) of variants that are common in ACpop (AF > 1%) but are not represented or rare (AF ≤ 1%) in SweGen. The average genome in ACpop consists of 4.20 M variants (3.98 M sites), where 3.44 M are SNPs and 0.77 M are indels, and carries 6,742 variants not found in dbSNP (v. 150).

**Table 2.**
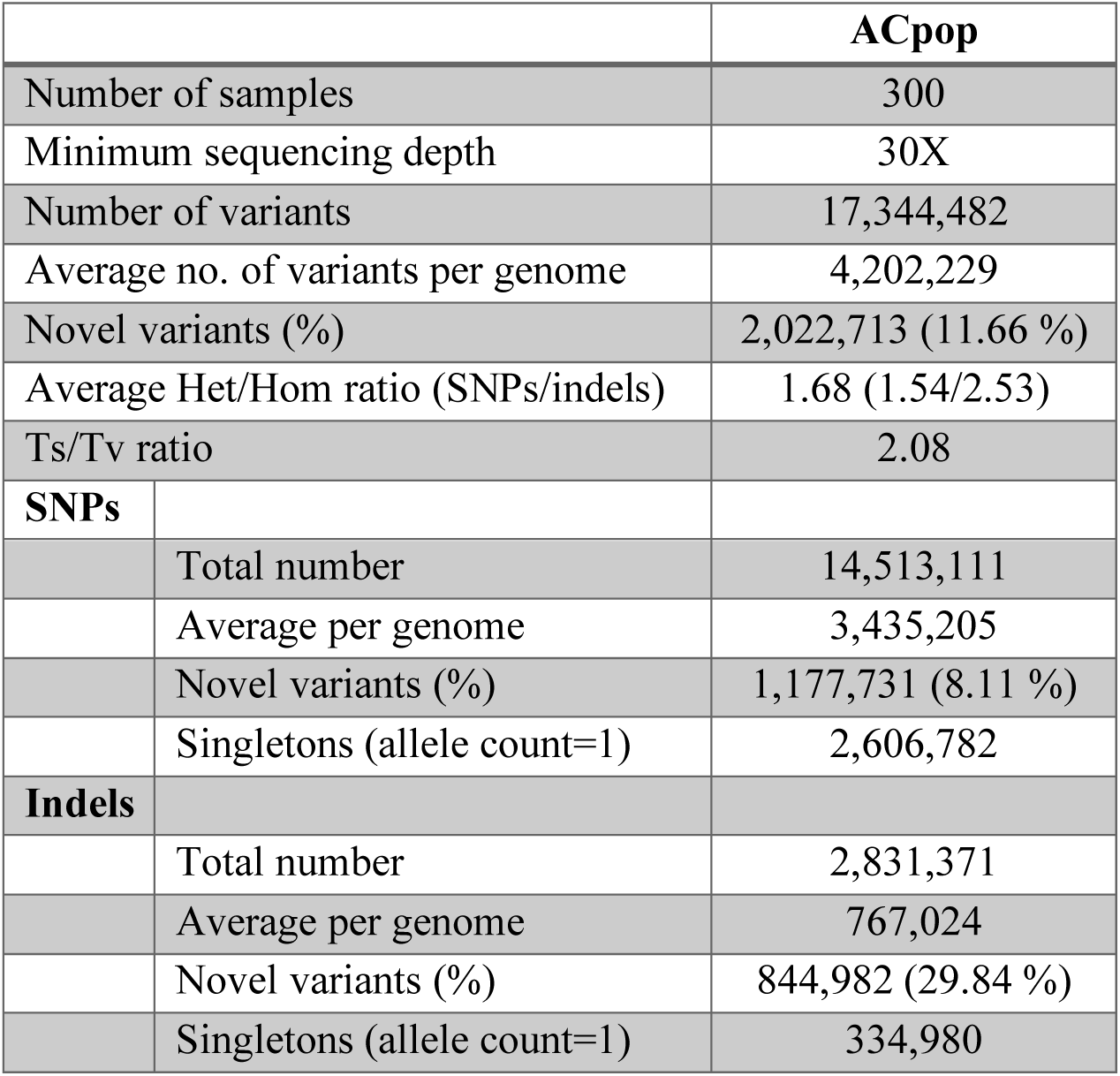
Summary table of the ACpop dataset. Novel variants is the number of variants in ACpop that are not present in the latest version of dbSNP (v. 150). Ts/Tv ratio is the ratio of SNP transitions and transversions, and Het/Hom ratio is the ratio between the number of heterozygous and homozygous occurrences of the variants.

|  |  | ACpop |
| --- | --- | --- |
| Number of samples |  | 300 |
| Minimum sequencing depth |  | 30X |
| Number of variants |  | 17,344,482 |
| Average no. of variants per genome |  | 4,202,229 |
| Novel variants (%) |  | 2,022,713 (11.66 %) |
| Average Het/Hom ratio (SNPs/indels) |  | 1.68 (1.54/2.53) |
| Ts/Tv ratio |  | 2.08 |
| **SNPs** |  |  |
|  | Total number | 14,513,111 |
|  | Average per genome | 3,435,205 |
|  | Novel variants (%) | 1,177,731 (8.11 %) |
|  | Singletons (allele count=1) | 2,606,782 |
| **Indels** |  |  |
|  | Total number | 2,831,371 |
|  | Average per genome | 767,024 |
|  | Novel variants (%) | 844,982 (29.84 %) |
|  | Singletons (allele count=1) | 334,980 |

### Subregional genetic structure of ACpop

In a global setting, the genetic variation of ACpop tends to show similarity with other European populations from the 1000g project, as evident by their co-localization within the first and second components of a joint PCA (Figure 2). To investigate the relationship between ACpop and other European populations in greater detail, a second PCA was performed with only the European subpopulations of the 1000g project. The ACpop samples displayed a distinct separation from members of the other populations in the first two principal components, indicating that there is genetic variation in ACpop that is not captured by sequencing efforts on other European populations (Figure 2B). The internal spread of ACpop in the first and second principal components is in part due to the fact that ACpop is a comparatively large sample in the context of 1000g project European populations. In addition, the pairwise F_ST_ between ACpop and the European populations was calculated. Of the five investigated European populations, ACpop appeared to be genetically closest to the British (GBR) population rather than the geographically neighboring Finnish (FIN) population (Table 3).

**Table 3.** Pairwise F_ST_ of European populations. The European populations are the subpopulations of the European super population of the 1000g project. The standard error of the corresponding F_ST_ value is given in parentheses.

|  | ACpop | GBR | CEU | IBS | TSI | FIN |
| --- | --- | --- | --- | --- | --- | --- |
| ACpop | -- | 0.004<br>(0.00006) | 0.004<br>(0.000055) | 0.008<br>(0.000097) | 0.009<br>(0.000176) | 0.006<br>(0.000076) |
| GBR |  | -- | 0.0<br>(0.000041) | 0.002<br>(0.000061) | 0.004<br>(0.000109) | 0.007<br>(0.000108) |
| CEU |  |  | -- | 0.002<br>(0.000073) | 0.004<br>(0.000134) | 0.006<br>(0.000106) |
| IBS |  |  |  | -- | 0.002<br>(0.000051) | 0.01<br>(0.000139) |
| TSI |  |  |  |  | -- | 0.012<br>(0.000192) |
| FIN |  |  |  |  |  | -- |

**Figure 2.**
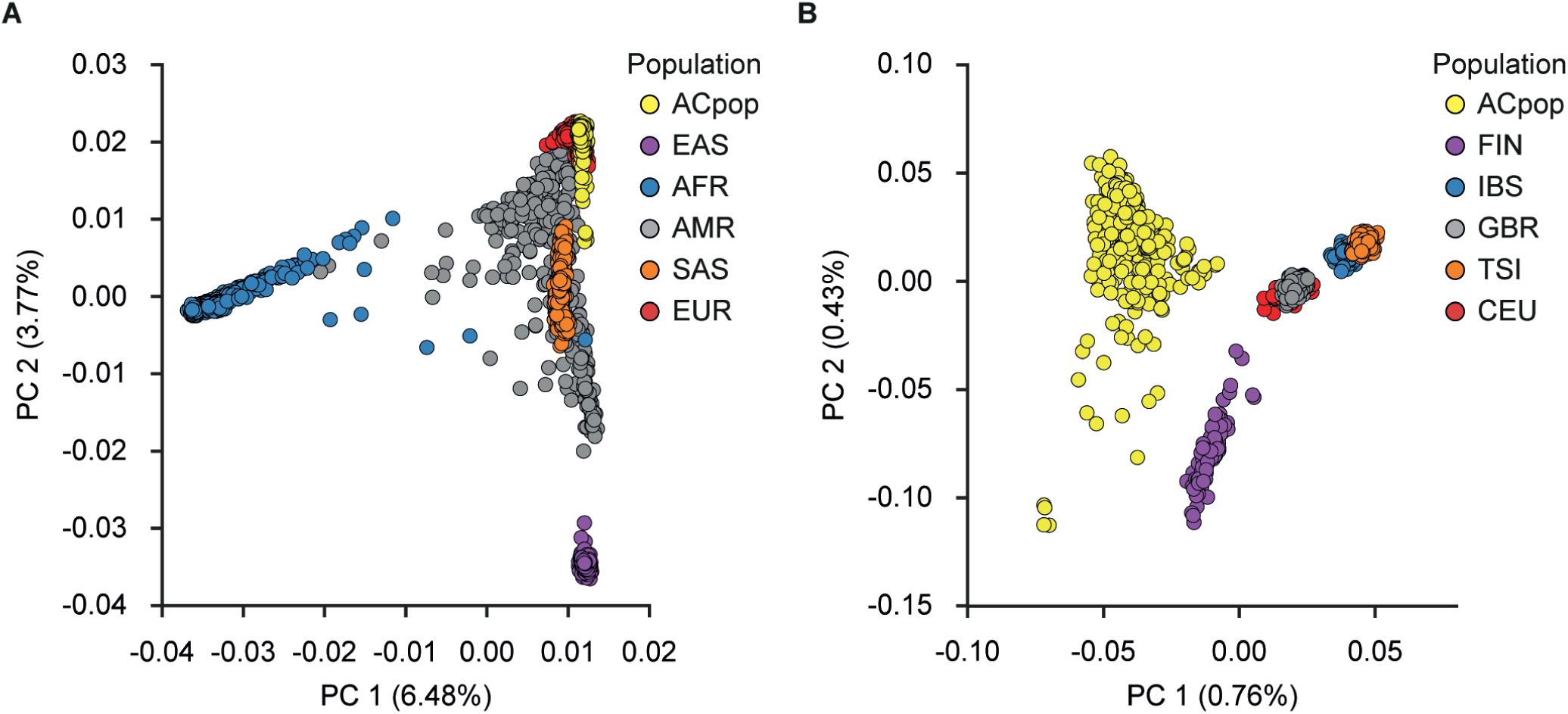
PCA of ACpop and the 1000g. Joint PCA of ACpop and the super populations (EUR: European, EAS: East Asian, AFR: African, AMR: Ad Mixed American, SAS: South Asian) of the 1000g project (**A**). Joint PCA of ACpop and the European populations (FIN: Finnish in Finland, CEU: Utah Residents (CEPH) with Northern and Western European Ancestry, GBR: British in England and Scotland, IBS: Iberian Population in Spain, TSI: Tuscany in Italy) of the 1000g project (**B**). The proportion of the variance explained by each principle component is indicated in the figure axis titles.

It was earlier demonstrated that there is a north-south gradient, where genetic subpopulations are located along the large river valleys, which extends from the mountains in the west to the coast in the east. The colonization of the inland and mountain regions with a relatively small founder population together with our contemporary analysis showing low mobility among individuals born in 1934 (Table 1), led us to ask if there is a genetic difference between the western and eastern populations. Västerbotten County can be divided into three geographical regions, a mountainous region, an inland region, and a coastal region (Figure 3). To further investigate any genetic structure within the county, a PCA was performed with the ACpop samples. By dividing the samples into groups corresponding to the three geographical regions, a subtle pattern emerged in which samples are placed along a gradient going from the coast, through the inland area and to the mountains (Figure 3). The same geographical division was used for calculating pairwise F_ST_. The intra-county genetic distance between the group of individuals associated with the mountainous region on the one hand, and the group of individuals associated with the coastal region on the other hand, is comparable to that of the genetic distance between the British (GBR) and Spanish (IBS) populations in the 1000g project (Table 3 and Table 4). The emerging structure within the county could be further stratified by the municipal association of the samples. Samples tend to cluster according to municipality in the first and second principal components in a way that mimics the geographical associations of the municipalities. That is to say, neighboring municipalities tend to be represented in the PCA by neighboring samples (Figure 4) which correlates with our demographic analysis (Table 1).

**Table 4.** Pairwise F_ST_ of Västerbotten subpopulations. The F_ST_ values were calculated pairwise between three geographically defined regions in Västerbotten County, the mountains, inland and coast. Parentheses contain the standard error of the corresponding F_ST_ value.

|  | Coast | Inland | Mountain |
| --- | --- | --- | --- |
| Coast | -- | 0.001<br>(0.000035) | 0.002<br>(0.000083) |
| Inland |  | -- | 0.001<br>(0.000067) |
| Mountain |  |  | -- |

**Figure 3.**
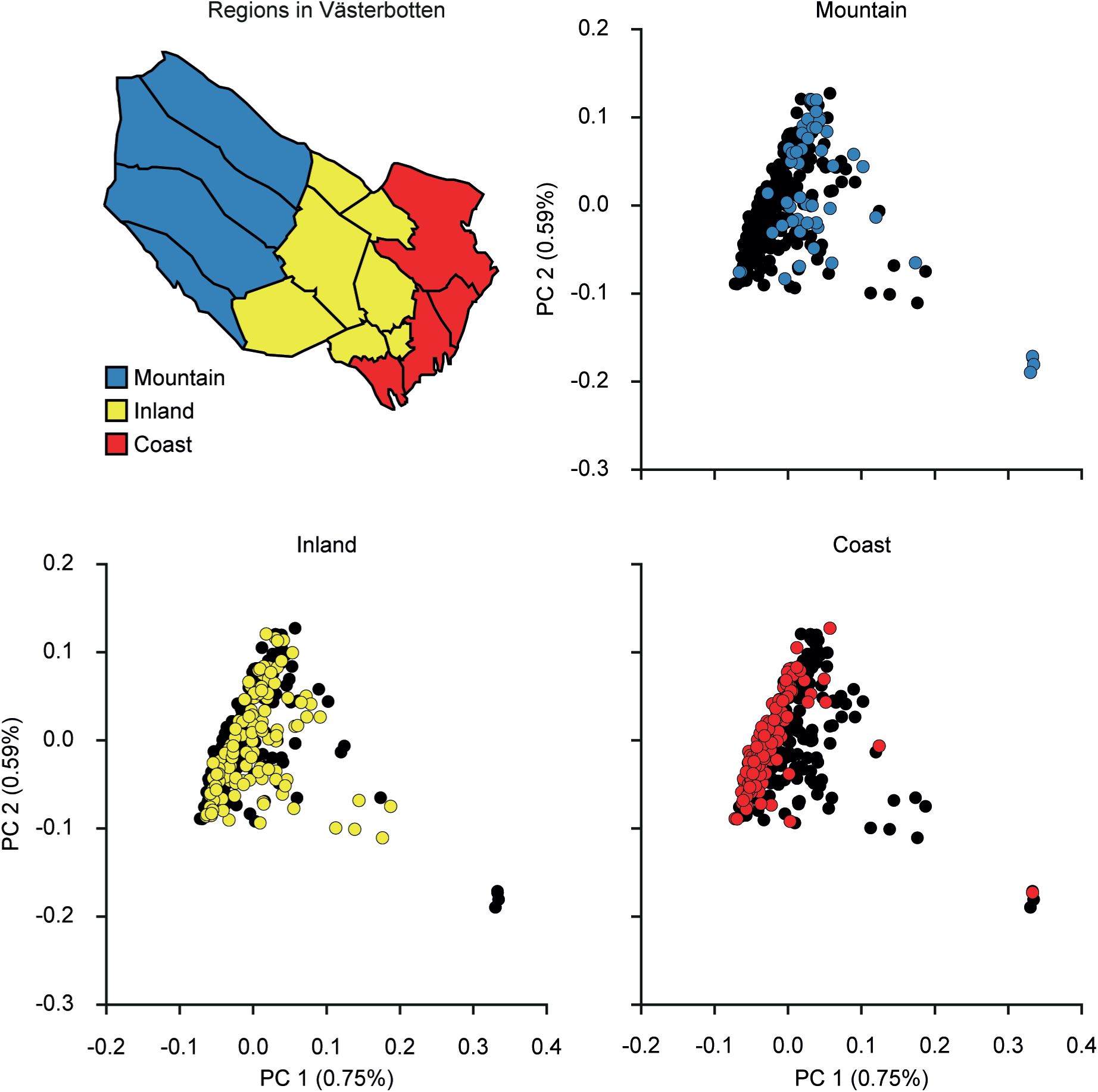
PCA of ACpop samples. The samples were plotted along principal components 1 and 2 and are colored according to three geographical regions in Västerbotten County. Top left panel: a map of Västerbotten County showing the 15 municipalities and the regional subdivision. The proportion of the variance explained by each principle component is indicated in the figure axis titles.

**Figure 4.**
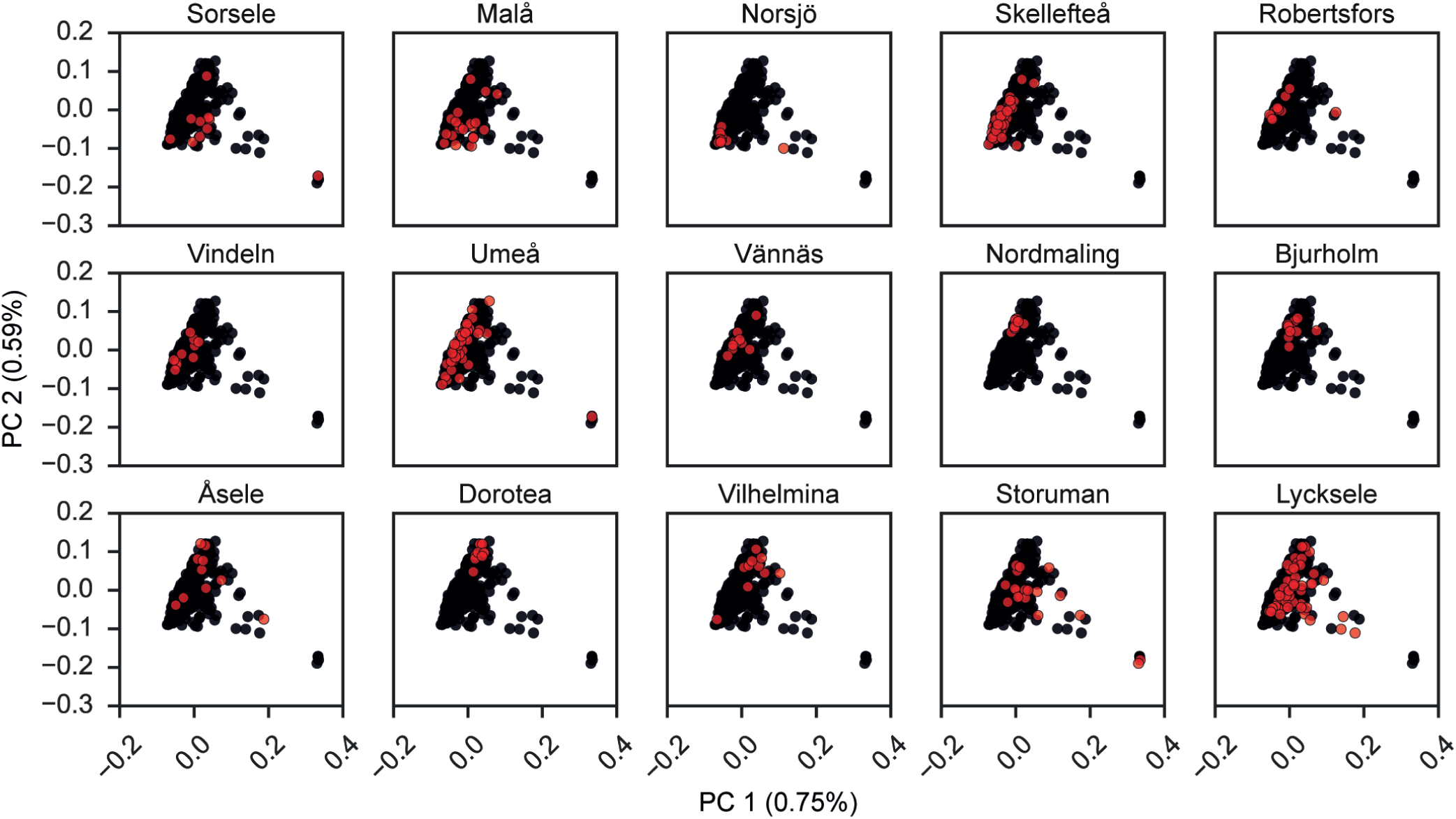
PCA of ACpop samples. The samples were plotted along the first and second components in 15 subplots, each corresponding to one of the municipalities in Västerbotten County. Samples associated with the municipality are colored red, and the rest are colored black. The proportion of the variance explained by each principle component is indicated in the figure axis titles.

Earlier array-based studies of the northern Swedish population have suggested an elevated level of autozygosity (14) and an increased number of homozygous segments (13). With access to a whole-genome sequenced, carefully selected and comparatively large population sample from a focused northern Swedish geographical area, we wanted to provide a better understanding of the existing homozygosity pattern. The degree of autozygosity can be estimated using the inbreeding coefficient *f* which represents the proportion of the genome that is homozygous by descent. The results point to a gradual increase of the average *f* from the coastal (*f=*0.014) region to the mountainous region (*f=*0.019) of Västerbotten County, with a total average *f* value of 0.015 (Table 4). The same gradual increase from coast to mountain can be witnessed for the proportion of individuals classified as showing signs of autozygosity, i.e. where *f* > 0.001 is satisfied (Supplementary Table 2).

### Effect of ACpop in imputation of national and regional cohorts

A whole genome sequenced reference population can be used as a reference panel for imputation in genome-wide association studies (GWAS). Previous studies have shown that population-specific reference panels can improve imputation accuracy in matching cohorts (41,42). To investigate the imputation performance of ACpop, we compared the number of variants on chromosome 20 that were imputed with a high *info* score, which is given by the imputation software as a measure of confidence of imputation of individual variants, when using (i) the 1000g reference panel, (ii) the ACpop reference panel, and (iii) a combination of the two reference panels. The number of variants imputed with high confidence (info ≥ 0.8) increased by 37% in a cohort from Västerbotten County (Figure 5A) and by 11% in a Swedish national cohort (Figure 5B) when using the combination of the reference panels, compared to using only the 1000g reference panel. For variants with minor allele frequency ≤ 5% the corresponding increase was 81% and 23% for the Västerbotten and Swedish cohort, respectively (Figure 5). For variants that were directly genotyped by array technology in the Västerbotten County and Swedish national cohorts, accuracy of imputation was assessed by means of r^2^ values (i.e. the correlation between the genotyped and imputed values for a particular variant). Accuracy of imputation of genotyped variants increased with 3.4% in the Västerbotten county cohort (mean r^2^=0.98) and 1.1% in the Swedish cohort (mean r^2^=0.95) when using the combination of 1000g and ACpop, compared to using the 1000g panel alone (Supplementary figure 3).

**Figure 5.**
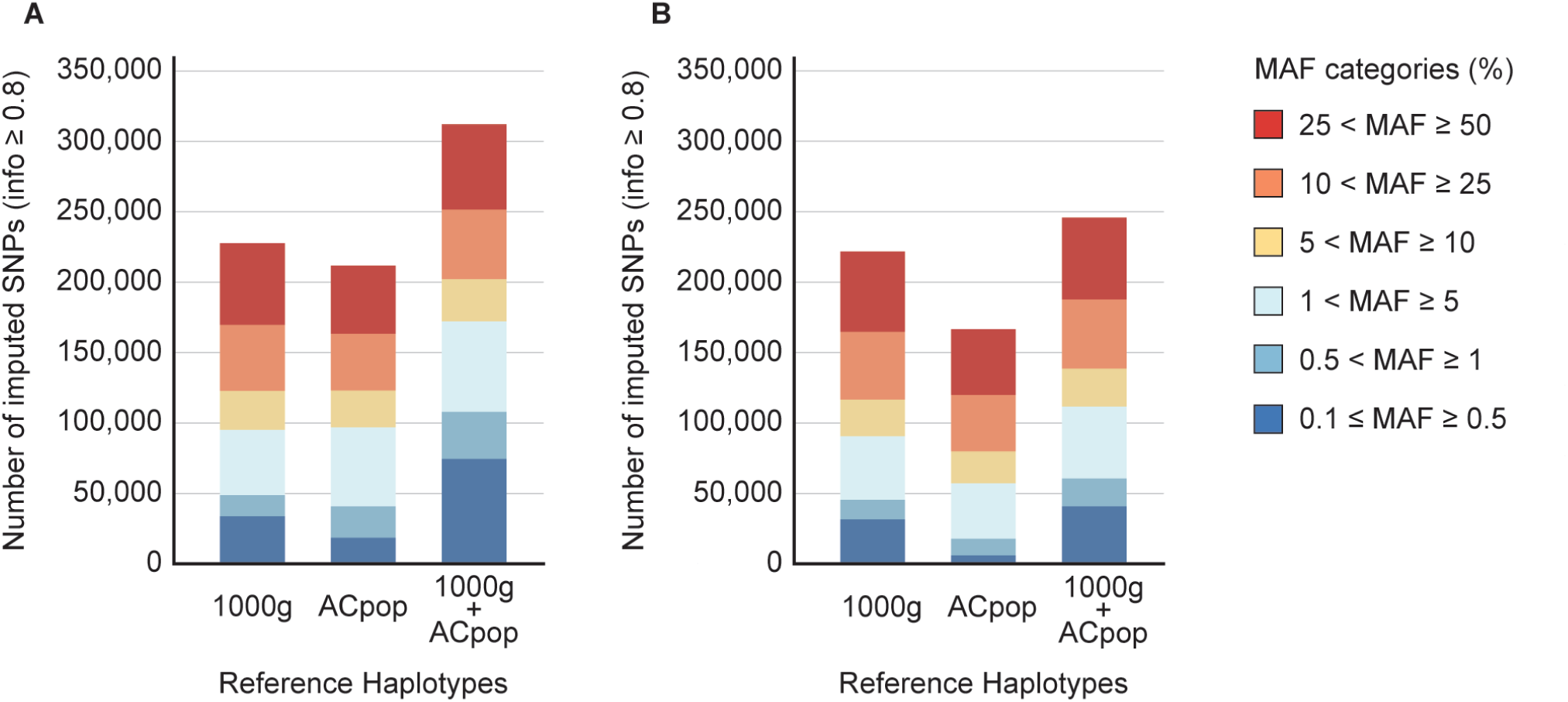
Imputation result. Unknown genotypes on chromosome 20 were imputed using either only the 1000g reference panel, only the ACpop reference panel, or using a combined reference panel from both panels. The number of high quality SNPs (info ≥ 0.8), aggregated by minor allele frequency (MAF), that were imputed in a Västerbotten cohort (A) and a Swedish national cohort (B) is indicated.

## Discussion

In this study, we have presented and characterized ACpop, a whole genome sequenced Västerbotten County population sample of 300 individuals which corresponds to 0.11% of the population. The selection of ACpop aimed in part to include as much as possible of the genetic diversity in Västerbotten County. Because geographic origin is a determining factor of genetic variation, a main component in the selection was to ensure a geographical spread across the county. In addition, using statistical design we selected diverse individuals with respect to health and life style associated variables in the biobank. Demographic investigations indicate that the population we selected from was, to a large extent, living in the same municipality in which they were born. Therefore, by basing the selection on the municipal association of the samples, we are confident that ACpop is representative of the genetic variation in Västerbotten. Furthermore, ACpop includes a large number of rare local variants that were not represented in dbSNP or SweGen. In all, ACpop is a resource that provides an unprecedented level of detail of the genetic landscape of Västerbotten County.

Previous studies describing the Swedish genetic landscape have highlighted genetic differences between northern Sweden and both southern Sweden (15) and other European populations (13,14). ACpop demonstrates in this study that not only is there a genetic substructure within Sweden but also within Västerbotten county. This is in line with indications of an elevated positive correlation of genetic and geographic distances demonstrated in a previous array-based study (14). The intra-county genetic distances between the mountain and coast regions as measured by F_ST_ are comparable to differences between major European populations in the 1000g project (Table 4). This substructure is also supported in the principal component analyses where the two largest components reflect the municipal and regional associations of the samples, respectively (Figure 3, Figure 4). In fact, our results suggest that association studies in Västerbotten might show an improvement if matched controls are taken not only from Västerbotten, but from the same municipality as each affected individual. Similar conclusions were drawn from studies on the Icelandic population with a genealogy that show some similarities to Västerbotten (43). Iceland was colonized by 8,000-16,000 settlers 1,100 years ago, maintained a population size of about 50,000 around 1850, followed by an expansion to its current size of about 300,000. This resulted in a population with genetic substructures even if the population was assumed to be homogeneous. Indication of the Västerbotten substructure is also given by the inbreeding analysis where the coefficient of inbreeding increased from the coast to the mountains. The comparatively high coefficient of inbreeding might be the result of a small gene pool associated with the small number of founders, in combination with population isolation. In addition, one study (44) that was limited to one coastal municipality suggested an increased occurrence of consanguineous marriages during the 19^th^ century which, if this is representative for the entire county, will influence the inbreeding coefficient of our samples.

Imputation is a method that is often used to increase the number of genetic variants that can be investigated in genetic association studies, and the 1000g is often the reference panel of choice for these analyses. But although the 1000g includes several genetically diverse populations, we found that including ACpop (n=300) together with the 1000g reference panel (n=2,504), had a positive impact on imputation performance in Västerbotten samples. The increase of confidently imputed variants was highest for variants with minor allele frequency ≤ 5%. Rare variants are underrepresented on the array and in our analyses of imputation accuracy. However, previous studies have shown that the addition of a genetically well-match reference panel increase imputation accuracy also for rare variants (42). The increase in imputation performance was not as striking in a Swedish national cohort, which again is likely to be the result of the genetic differences between the north and south of Sweden. Imputation performance is expected to increase as haplotypes are added to the imputation reference panel (45). However, the modest increase in imputation performance in the Swedish national cohort suggests that the distinct increase in imputation performance in the Västerbotten cohort is, to a large extent, due to the addition of reference samples that are genetically well-matched. Our results suggest that association studies in populations that are not well represented by panels such as 1000g would greatly benefit by the addition of reference subjects for imputation who are drawn from the same geographical region as the studied individuals.

## Acknowledgments

We are grateful to all participants of the VIP cohort of the Northern Sweden Health and Disease Study (NSHDS), the staff at the Department of Biobank Research, Umeå University. Illumina sequencing was performed by the National Genomics Infrastructure (NGI), which is hosted by SciLifeLab in Stockholm and Uppsala. The computations were performed using resources provided by the Swedish National Infrastructure for Computing (SNIC) through the Uppsala Multidisciplinary Center for Advanced Computational Science (UPPMAX). P.I.O. was supported by a grant from the Knut and Alice Wallenberg Foundation to the Wallenberg Advanced Bioinformatics Infrastructure. This work was funded by a project grant (2011.0042) from the Knut and Alice Wallenberg Foundation (E.J., B.S.M., J.T.).

## Notes

**Grants:** This work was funded by a project grant (2011.0042) from the Knut and Alice Wallenberg Foundation (E.J., B.S.M., J.T.).

**Conflict of interest:** The authors declare no conflict of interest.

https://swefreq.nbis.se/dataset/ACpop

## References

1. Auton A, Abecasis GR, Altshuler DM, et al. A global reference for human genetic variation. Nature. 2015 Sep 30;526(7571):68–74.

2. Lek M, Karczewski KJ, Minikel E V., et al. Analysis of protein-coding genetic variation in 60,706 humans. Nature. 2016 Aug 17;536(7616):285–91.

3. Altshuler DM, Gibbs RA, Peltonen L, et al. Integrating common and rare genetic variation in diverse human populations. Nature. 2010 Sep 2;467(7311):52–8.

4. Coventry A, Bull-Otterson LM, Liu X, et al. Deep resequencing reveals excess rare recent variants consistent with explosive population growth. Nat Commun. 2010 Nov 30;1(8):131–6.

5. Keinan A, Clark AG. Recent Explosive Human Population Growth Has Resulted in an Excess of Rare Genetic Variants. Science (80-). 2012;336:740–3.

6. Fu W, O’Connor TD, Jun G, et al. Analysis of 6,515 exomes reveals the recent origin of most human protein-coding variants. Nature. 2012 Nov 28;493(7431):216–20.

7. Nelson MR, Wegmann D, Ehm MG, et al. An Abundance of Rare Functional Variants in 202 Drug Target Genes Sequenced in 14,002 People. Science (80-). 2012 Jul 6;337(6090):100–4.

8. Gravel S, Henn BM, Gutenkunst RN, et al. Demographic history and rare allele sharing among human populations. Proc Natl Acad Sci. 2011;108(29):11983–8.

9. The Genome of the Netherlands Consortium. Whole-genome sequence variation, population structure and demographic history of the Dutch population. Nat Genet. 2014;46(8):818–25.

10. Gudbjartsson DF, Helgason H, Gudjonsson S a, et al. Large-scale whole-genome sequencing of the Icelandic population. Nat Genet. 2015 Mar 25;47(5):435–44.

11. Nagasaki M, Yasuda J, Katsuoka F, et al. Rare variant discovery by deep whole-genome sequencing of 1,070 Japanese individuals. Nat Commun. 2015;6:1–13.

12. Ameur A, Dahlberg J, Olason P, et al. SweGen: a whole-genome data resource of genetic variability in a cross-section of the Swedish population. Eur J Hum Genet. 2017 Nov 23;25(11):1253–60.

13. Humphreys K, Grankvist A, Leu M, et al. The genetic structure of the Swedish population. PLoS One. 2011;6(8):e22547.

14. Salmela E, Lappalainen T, Liu J, et al. Swedish population substructure revealed by genome-wide single nucleotide polymorphism data. PLoS One. 2011 Jan 9;6(2):e16747.

15. Karlsson AO, Wallerström T, Götherström A, Holmlund G. Y-chromosome diversity in Sweden – A long-time perspective. Eur J Hum Genet. 2006 Aug 24;14(8):963–70.

16. Statistics Sweden. Land and water area in square kilometre by region, type of area and year [Internet]. 2017 [cited 2018 Jan 19]. Available from: http://www.statistikdatabasen.scb.se/pxweb/en/ssd/START__MI__MI0802/Areal2012/?rxid=6ad4df53-3608-4fc9-b2af-faab9be4a126

17. Statistics Sweden. Population 1 November by region, age and sex. Year 2002 - 2017 [Internet]. 2017 [cited 2018 Jan 19]. Available from: http://www.statistikdatabasen.scb.se/pxweb/en/ssd/START__BE__BE0101__BE0101A/FolkmangdNov/?rxid=4d6d2f0f-c0fc-4ab6-8cd9-77032b5e2ee2

18. Palm LA. Folkmängden i Sveriges socknar och kommuner 1571-1997 : med särskild hänsyn till perioden 1571-1751. Göteborg: L. A. Palm; 2000. 199 p.

19. Bylund E. Koloniseringen av Botniaregionen. In: Edlund L-E, Beckman L, editors. Botnia : en nordsvensk region. Höganäs: Bra böcker; 1994. p. 86–98.

20. Sköld P, Axelsson P. The northern population development; colonization and mortality in Swedish Sápmi, 1776-1895. Int J Circumpolar Health. 2008;67(1):27–42.

21. Einarsdottir E, Egerbladh I, Beckman L, Holmberg D, Escher SA. The genetic population structure of northern Sweden and its implications for mapping genetic diseases. Hereditas. 2007;144(5):171–80.

22. Norberg M, Blomstedt Y, Lönnberg G, et al. Community participation and sustainability – evidence over 25 years in the Västerbotten Intervention Programme. Glob Health Action. 2012 Dec 17;5(1):19166.

23. DePristo MA, Banks E, Poplin R, et al. A framework for variation discovery and genotyping using next-generation DNA sequencing data. Nat Genet. 2011 May;43(5):491–8.

24. Van der Auwera GA, Carneiro MO, Hartl C, et al. From FastQ data to high confidence variant calls: the Genome Analysis Toolkit best practices pipeline. Curr Protoc Bioinforma. 2013;43(1110):11.10.1-33.

25. Li H, Durbin R. Fast and accurate short read alignment with Burrows-Wheeler transform. Bioinformatics. 2009 Jul 15;25(14):1754–60.

26. Picard [Internet]. Available from: http://broadinstitute.github.io/picard

27. McKenna A, Hanna M, Banks E, et al. The Genome Analysis Toolkit: a MapReduce framework for analyzing next-generation DNA sequencing data. Genome Res. 2010 Sep;20(9):1297–303.

28. Okonechnikov K, Conesa A, García-Alcalde F. Qualimap 2: advanced multi-sample quality control for high-throughput sequencing data. Bioinformatics. 2015 Oct 1;32(2):btv566.

29. Manichaikul A, Mychaleckyj JC, Rich SS, Daly K, Sale M, Chen W. Robust relationship inference in genome-wide association studies. 2010;26(22):2867–73.

30. Patterson N, Price AL, Reich D. Population structure and eigenanalysis. PLoS Genet. 2006;2(12):2074–93.

31. Chang CC, Chow CC, Tellier LC, Vattikuti S, Purcell SM, Lee JJ. Second-generation PLINK: rising to the challenge of larger and richer datasets. Gigascience. 2015;4:7.

32. Rajaraman P, Melin BS, Wang Z, et al. Genome-wide association study of glioma and meta-analysis. Hum Genet. 2012 Dec 11;131(12):1877–88.

33. Delaneau O, Marchini J, Zagury J-F. A linear complexity phasing method for thousands of genomes. Nat Methods. 2011 Dec 4;9(2):179–81.

34. Howie BN, Donnelly P, Marchini J. A flexible and accurate genotype imputation method for the next generation of genome-wide association studies. PLoS Genet. 2009 Jun;5(6):e1000529.

35. Howie B, Fuchsberger C, Stephens M, Marchini J, Abecasis GR. Fast and accurate genotype imputation in genome-wide association studies through pre-phasing. Nat Genet. 2012 Jul 22;44(8):955–9.

36. Delaneau O, Zagury J-F, Marchini J. Improved whole-chromosome phasing for disease and population genetic studies. Nat Methods. 2013;10(1):5–6.

37. Gazal S, Sahbatou M, Babron MC, Génin E, Leutenegger AL. FSuite: Exploiting inbreeding in dense SNP chip and exome data. Bioinformatics. 2014;30(13):1940–1.

38. Gazal S, Sahbatou M, Babron M-C, Génin E, Leutenegger A-L. High level of inbreeding in final phase of 1000 Genomes Project. 2015;

39. Graffelman J, Moreno V. The mid p-value in exact tests for Hardy-Weinberg equilibrium. Stat Appl Genet Mol Biol. 2013 Jan 1;12(4):433–48.

40. Leutenegger AL, Sahbatou M, Gazal S, Cann H, Génin E. Consanguinity around the world: What do the genomic data of the HGDP-CEPH diversity panel tell us? Eur J Hum Genet. 2011;19(5):583–7.

41. Surakka I, Kristiansson K, Anttila V, et al. Founder population-specific HapMap panel increases power in GWA studies through improved imputation accuracy and CNV tagging. Genome Res. 2010 Oct;20(10):1344–51.

42. Mitt M, Kals M, Pärn K, et al. Improved imputation accuracy of rare and low-frequency variants using population-specific high-coverage WGS-based imputation reference panel. Eur J Hum Genet. 2017;2551(10):869–76.

43. Helgason A, Yngvadóttir B, Hrafnkelsson B, Gulcher J, Stefánsson K. An Icelandic example of the impact of population structure on association studies. Nat Genet. 2005 Jan 19;37(1):90–5.

44. Bittles AH, Egerbladh I. The influence of past endogamy and consanguinity on genetic disorders in northern Sweden. Ann Hum Genet. 2005;69(5):549–58.

45. Howie B, Marchini J, Stephens M. Genotype Imputation with Thousands of Genomes. G3; Genes|Genomes|Genetics. 2011 Nov 4;1(6):457–70.

