## Supplemental Table 1 for "A whole-genome sequenced control population in northern Sweden reveals subregional genetic differences"

*Supplementary Table 1* – **Variables used in the selection.**

| **Variable** |
|---|
| Age |
| Height |
| Weight |
| BMI |
| Total cholesterol |
| Hdl cholesterol |
| Triglycerides |
| Blood glucose, fasting 0 hours |
| Blood glucose 2 hours |
| Systolic blood pressure |
| Diastolic blood pressure |
| Educational level |
| Long-term sickness (6 months) |
| Self-estimated health status compared to others of same age |
| Parent/sibling cerebral hemorrhage/thrombosis or cardiac infarction before age 60 |
| Parent/sibling have diabetes |
| Informed of high blood pressure at any time |
| Diabetes |
| Heart attack that lead to hospitalization |
| Physically heavy job |
| Mentally demanding job |
| Physical activity before age of 20 |
| Teetotaler |
| Smoker |
| Smoking start age |
| Snuffer |
| Physical activity index |
