## Supplemental Table 2 for "A whole-genome sequenced control population in northern Sweden reveals subregional genetic differences"

*Supplementary Table 2* - **Summary of inbreeding coefficient estimation.** The results are presented in total and divided into three geographic regions. The inbreeding coefficient f can be considered to be the ratio of the genome of an individual that is homozygous by descent. The average f value is the average taken across the individuals in each category. The inbred status of a sample is the result of testing if f is significantly different from 0.001 using a likelihood test.

| | Mountains | Inland | Coast | Total |
|---:|---:|---:|---:|---:|
| No. samples | 50 | 120 | 130 | 300 |
| Average f | 0.019 | 0.015 | 0.014 | 0.015 |
| No. samples classified as inbred | 36 | 80 | 85 | 201 |
| Samples classified as inbred (%) | 72% | 66.7% | 65.4% | 67% |
