## Supplemental Figure 1 for "A whole-genome sequenced control population in northern Sweden reveals subregional genetic differences"

*Supplementary Figure 1*

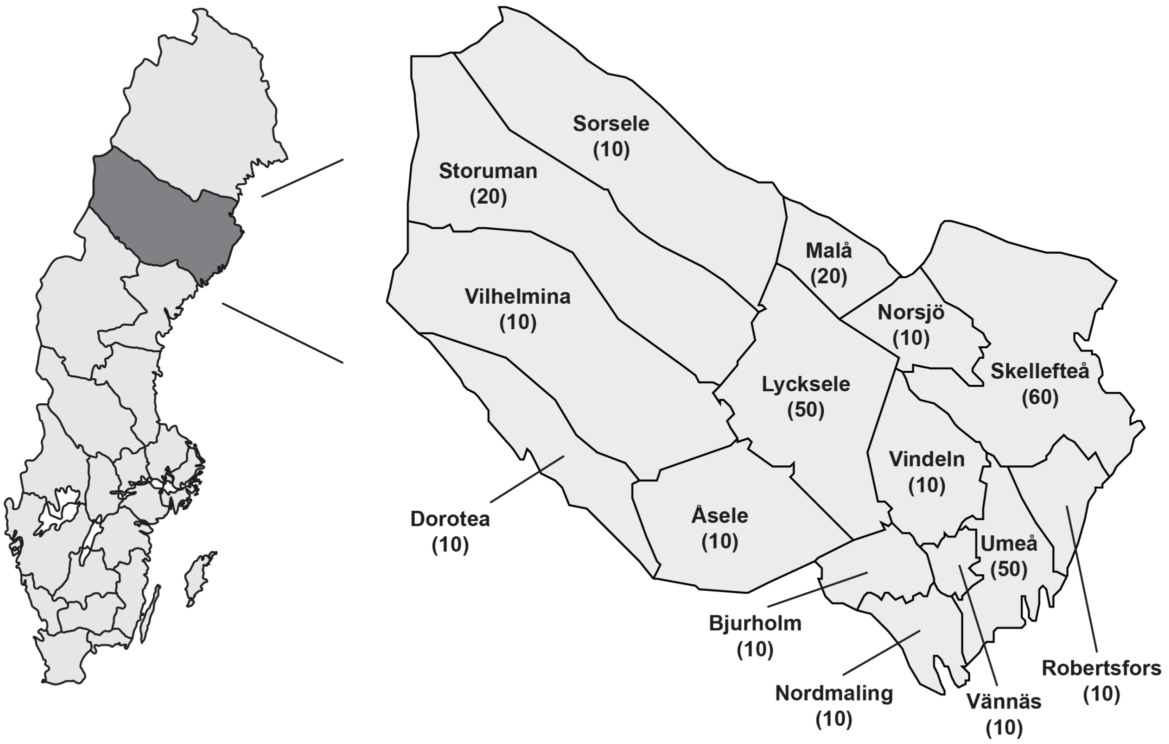


*Supplementary figure 1* – **Geographical overview of Västerbotten County.** The location of Västerbotten County in Sweden (grey) (left panel). A map of Västerbotten County with the location of all 15 municipalities and their names (right panel). The number of samples drawn from each municipality is indicated within parentheses below the name of the corresponding municipality.
