## Supplemental Figure 2 for "A whole-genome sequenced control population in northern Sweden reveals subregional genetic differences"

Supplementary Figure 2

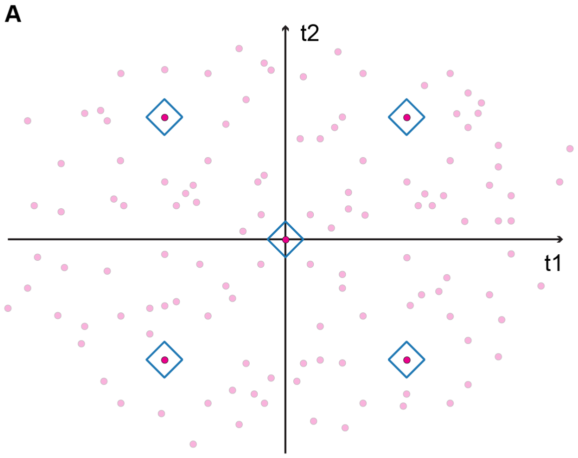


*Supplementary figure 2* - **Designs used for selection of samples for ACpop.** The figure illustrates a schematic score plot of the first and second component from a PCA model. Pink dots represent the data points (samples) and diamonds indicate selected samples. The baseline selection for all municipalities was made according to a full factorial design (A), whereas for the more populous municipalities an extended design was used (B).
