## Supplemental Figure 3 for "A whole-genome sequenced control population in northern Sweden reveals subregional genetic differences"

Supplementary Figure 3

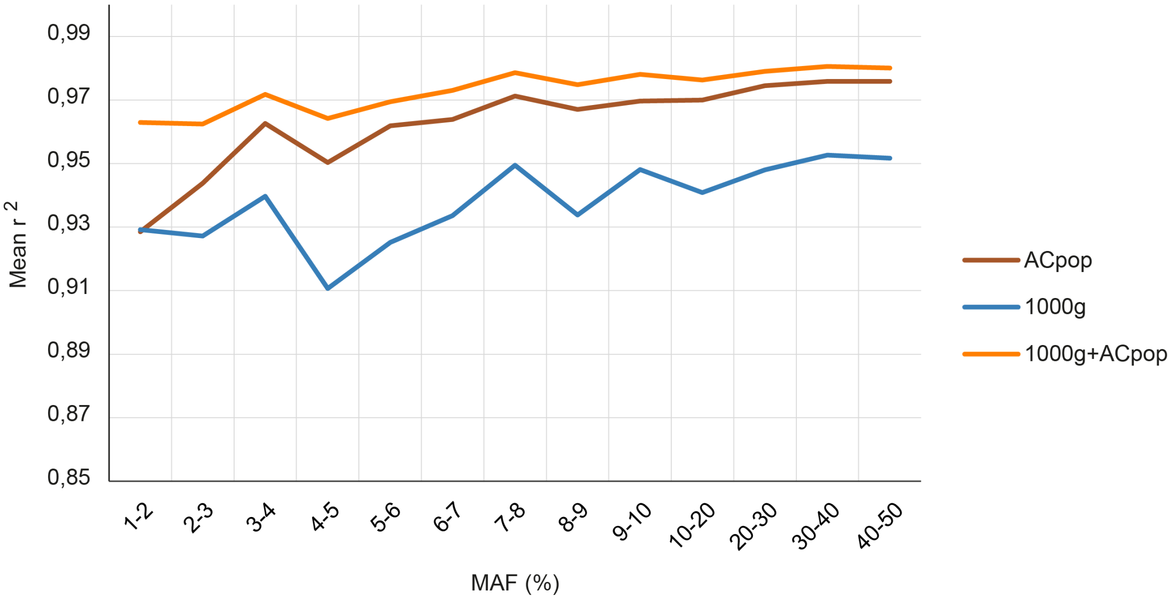


*Supplementary figure 3* – **Imputation accuracy.** The mean $r^2$ value for imputed and genotyped variants when using 1) only ACpop, 2) only 1000g, and 3) both ACpop and 1000g as the reference panel. The results are binned according to the expected minor allele frequency (MAF).
